# Biological capacities clearly define a major subdivision in Domain Bacteria

**DOI:** 10.1101/335083

**Authors:** Raphaël Méheust, David Burstein, Cindy J. Castelle, Jillian F. Banfield

## Abstract

Phylogenetic analyses separate candidate phyla radiation (CPR) bacteria from other bacteria, but the degree to which their proteomes are distinct remains unclear. Here, we leveraged a proteome database that includes sequences from thousands of uncultivated organisms to identify protein families and examine their organismal distributions. We focused on widely distributed protein families that co-occur in genomes, as they likely foundational for metabolism. Clustering of genomes using the protein family presence/absence patterns broadly recapitulates the phylogenetic structure of the tree, suggesting persistence of core sets of protein families after lineage divergence. CPR bacteria group together and away from all other bacteria and archaea, in part due to novel proteins, some of which may be involved in cell-cell interactions. The diversity of combinations of protein families in CPR may exceed that of all other bacteria. Overall, the results extend the phylogeny-based suggestion that the CPR represent a major subdivision within Bacteria.

## Introduction

Metagenomic investigations of microbial communities have generated genomes for a huge diversity of bacteria and archaea, many from little studied or previously unknown phyla (Castelle and Banfield, 2018). For example, a study of an aquifer near the town of Rifle, Colorado generated 49 draft genomes for several groups of bacteria, some of which were previously known only based on 16S rRNA gene surveys and others that were previously unknown (Wrighton et al., 2012). Draft genomes for bacteria from related lineages were obtained in a single cell sequencing study that targeted samples from a broader variety of environment types (Rinke et al., 2013). Based on the consistently small predicted genome sizes for bacteria from these groups, groundwater filtration experiments targeting ultra-small organisms were conducted to provide cells for imaging (Luef et al., 2015) and DNA for increased genomic sampling. The approach yielded almost 800 genomes from a remarkable variety of lineages that place together phylogenetically. This monophyletic group were described as the Candidate Phyla Radiation (CPR) (Brown et al., 2015). CPR bacterial genomes have since been recovered from the human microbiome (He et al., 2015), drinking water (Danczak et al., 2017), marine sediment (Orsi et al., 2017), deep subsurface sediments (Anantharaman et al., 2016), soil (Starr et al., 2017), the dolphin mouth (Dudek et al., 2017) and other environments. Thus, it appears that CPR bacteria are both hugely diverse and widespread across earth’s environments.

Metabolic analyses of CPR genomes consistently highlight major deficits in biosynthetic potential, leading to the prediction that most of these bacteria live as symbionts. Cultivation from human oral samples highlighted the attachment of a CPR member of the lineage Saccharibacteria (TM7) to the surface of an *Actinomyces odontolyticus* bacteria (He et al., 2015). Another episymbiotic association has been described between a CPR organism from the Parcubacteria superphylum and an eukaryotic host (Gong et al., 2014). However, most CPR organisms are likely symbionts of bacteria or archaea, given their abundance and diversity in samples that have few, if any, eukaryotes (Castelle and Banfield, 2018).

When first described, the CPR was suggested to comprise at least 15% of all Bacteria (Brown et al., 2015). Subsequently, Hug et al. placed a larger group of CPR genome sequences in context via construction of a three domain tree and noted that the CPR could comprise as much as 50% of all bacterial diversity (Hug et al., 2016). The CPR placed as the basal group in the bacterial domain in a concatenated ribosomal protein tree, but the deep branch positions were not sufficiently well supported to enable a conclusion regarding the point of divergence of CPR from other bacteria. The scale of the CPR is also controversial. For example, Parks et al. suggested that the group comprises no more than 26.3% of bacterial phylum-level lineages (Parks et al., 2017). Their analyses were based on a FastTree (Price et al., 2009) constructed using an alignment of 120 concatenated proteins, some of which do not occur in CPR. Removal of genomes with < 40% of the alignment length resulted in exclusion of most of the CPR sequences. Regardless, the CPR clearly represents a huge segment of bacterial diversity.

To date, most studies have predicted CPR metabolic traits using one or a few genomes. Lacking are studies that look radiation-wide at the distribution of capacities that are widespread thus likely contribute core functions, including those encoded by hypothetical proteins. Moreover, examination of genetic potential across the CPR and general comparisons of CPR and non-CPR bacteria have been very limited. Here, we leveraged a large set of publicly available, high quality genomes of CPR and non-CPR bacteria to address these questions. We clustered protein sequences from 3,598 genomes into families and evaluated the distribution of these protein families over genomes. By focusing only on protein families that are common in CPR bacteria and/or non-CPR bacteria, we demonstrate a major subdivision within the bacterial domain without reliance on gene or protein sequence phylogenies.

## Results

### Clustering of proteins into families and assessment of cluster quality

We collected 3,598 genomes from four published datasets (Anantharaman et al., 2016; Brown et al., 2015; Castelle et al., 2015; Probst et al., 2017). The dataset includes 2,321 CPR genomes from 65 distinct phyla (1,953,651 proteins), 1,198 non-CPR bacterial genomes from 50 distinct phyla (3,018,597 proteins) and 79 archaeal genomes (89,709 proteins) (Figure 1). Note that this huge sampling of Candidate Phyla was only possible due to genomes reconstructed in the last few years (Figure 1). We clustered the 5,061,957 protein sequences in a multi-step procedure (see materials and methods) to generate groups of homologous proteins. The objective was to convert amino acid sequences into units of a common language, allowing us to compare the proteomes across a huge diversity of genomes. This resulted in 21,859 clusters that were present in at least 5 distinct genomes. These clusters are henceforth referred to as protein families.

**Figure 1.**
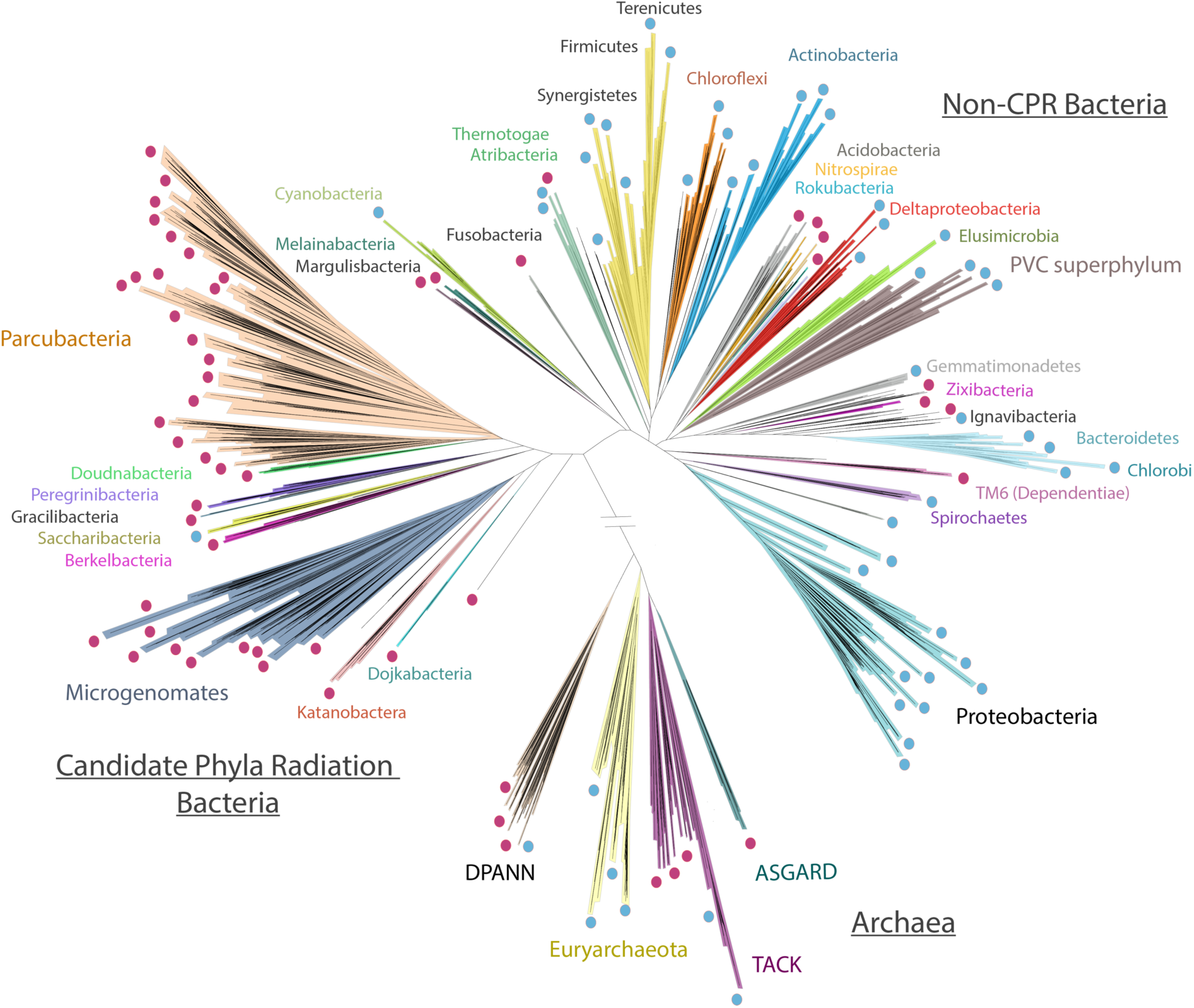
Schematic tree illustrating the phylogenetic sampling used in this study (the diagram is based on a tree published recently in (Castelle and Banfield, 2018)). Lineages that were included in the datasets are highlighted with a dot. Lineages lacking an isolated representative are highlighted with red dots. The number of genomes per lineages is available in Figure S1.

To assess the extent to which the protein clusters group together proteins with shared functions, we analyzed some families with well-known functions, such as the 16 ribosomal proteins that are commonly used in phylogeny. Because these proteins are highly conserved, we expect one protein family per ribosomal subunit. For instance, we expected to have all proteins annotated as the large subunit 3 (RPL3) be clustered into the same family. For 10 out 16 subunits, all proteins clustered into one single family (Table S1). The six remaining ribosomal subunits clustered into several families. However, one family always contained >95% of the proteins (Table S1). Interestingly, for five of the six subunits represented by more than one family, the fragmentation was a result of differences in protein length due to amino-acid extensions in the C-terminal or N-terminal regions. For instance, each of the 10 proteins from family 3.6k.fam10722 (consisting of RPL22 proteins, See Table S1) carries an extra 50 amino acids in the C-terminal region that is lacking in proteins of family 3.6k.fam00371 (Figure S1A). However, proteins from both families carry the domain of the large ribosomal subunit 22 (Figure S1A). We annotated our protein dataset using the KEGG annotations and systematically verified that the protein family groupings approximate functional annotations. For each KEGG accession, we reported the family which contains the highest ratio of proteins annotated with that KEGG accession. The distribution of the highest ratios shows that the majority of KEGG accessions is present in one major family (Figure S1B). We also checked the level of annotation admixture within the families. For each family, we computed the ratio of the KEGG accessions that are different than the most abundant accession (Figure S1C). The distribution of the ratios near 0 indicates that the vast majority of the families has no annotation admixture.

### The distribution of widespread proteins subdivides CPR from all other Bacteria

For definition of protein families, we chose a dataset that includes sequences from a huge diversity of uncultivated lineages and (unlike most reference genome datasets), genomes from the majority of all bacterial phyla (Figure 1). We constructed an array of the 3,598 genomes (rows) vs. all protein families (columns) and hierarchically clustered the genomes based on profiles of protein family presence/absence. The families were also hierarchically clustered based on profiles of genome presence/absence (Figure 2A). Notably, the distinct pattern of protein family presence/absence in CPR genomes separates them from almost all non-CPR bacteria and from archaea (Figure 2A).

**Figure 2.**
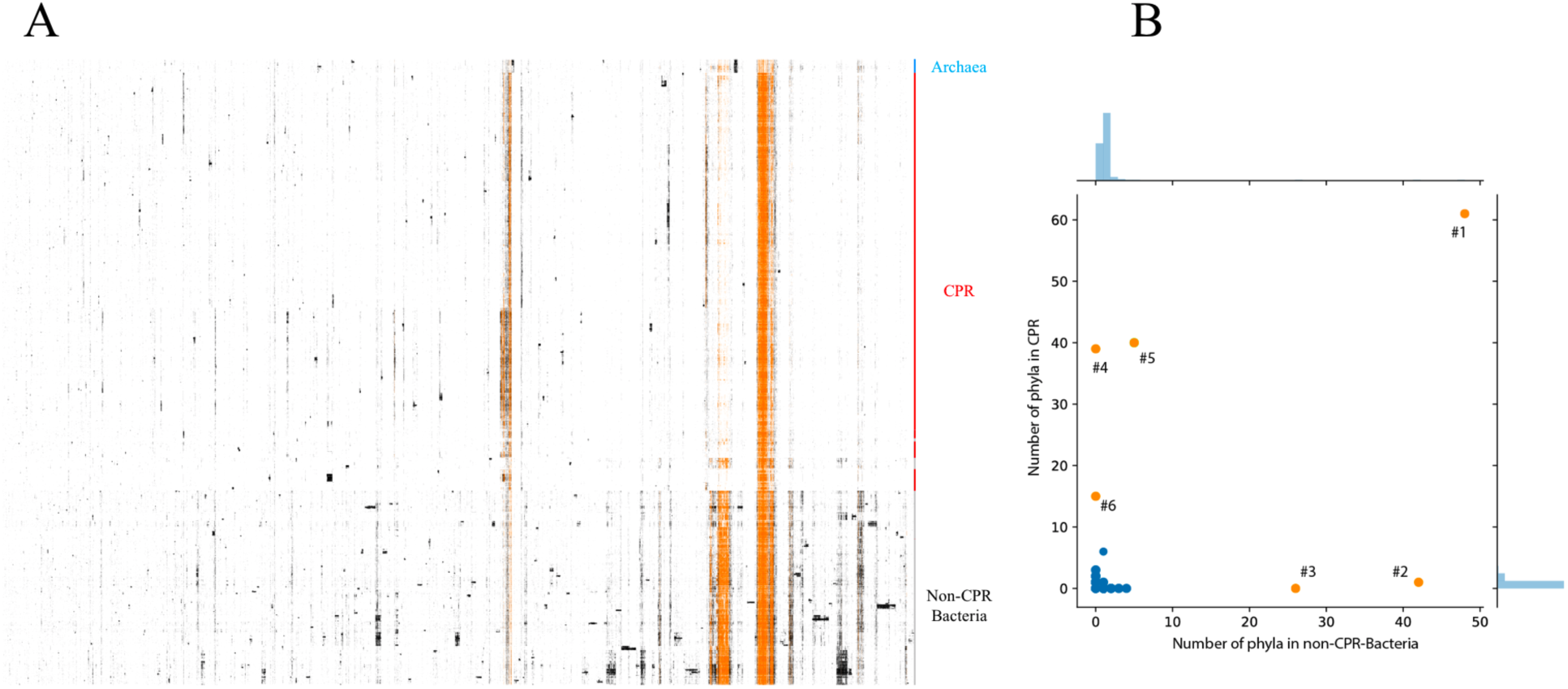
A. The distribution of the 21,859 families (columns) across the 3,598 prokaryotic genomes (rows). Data are clustered based on the presence (black) / absence (white) profiles (Jaccard distance, hierarchical clustering using a complete linkage) (Archaea: blue, CPR: red, non-CPR-Bacteria: gray). The patterns in orange correspond to the presence/absence patterns of the 786 widespread families that were retained for further analysis. B. The phyla distribution of the 156 modules of proteins in CPR (y-axis) and non-CPR-Bacteria (x-axis). Each dot corresponds to a module. The orange dots correspond to the 6 widespread modules that have been kept for further analysis.

Certain protein families cluster together due to co-existence in multiple genomes (blocks of black and orange dots in Figure 2A). Strikingly, some blocks with numerous families are widespread in non-CPR bacteria while mostly absent in CPR (Figure 2A), they may explain the observed separation of the CPR from the non-CPR Bacteria. We decided to focus on the large blocks of families that are widespread among the 115 bacterial phyla analyzed. We identified co-occurring blocks of protein families (sharing similar patterns of presence/absence across the genomes) using the Louvain algorithm (Blondel et al., 2008). We defined modules as blocks of co-occurring protein families containing at least 10 families. 7,962 protein families could be assigned to 156 modules. The remaining 13,997 protein families were not considered further. 150 of the 156 modules are sparsely distributed among the 115 bacterial phyla (blue dots in Figure 2B); these were also excluded from further analysis so that the study could focus only on the six modules that occur in most CPR bacterial genomes, in most non-CPR bacterial genomes or in both (orange dots in Figure 2B). Some modules also occur in archaeal genomes, so archaeal genomes were retained in the study.

One module, containing many core information system proteins, is essentially ubiquitous across the dataset (orange dot #1 in Figure 2B). Two modules are present in at least 10 non-CPR bacterial phyla (orange dots #2, 3 in Figure 2B). Strikingly, these modules are mostly absent in CPR bacteria. Three modules occur in more than 10 CPR bacterial phyla (orange dots #4, 5, 6 in Figure 2B).

The six numbered modules comprise 786 protein families that we consider to be widespread across the bacterial domain (blocks of orange dots in Figure 1.A). Unsurprisingly, given their widespread distribution, most these 786 families are involved in well-known functions, including replication, transcription and translation to basic metabolism (energy, nucleotides, amino-acids, cofactors and vitamins) and environmental interactions (membrane transport such as the Sec pathway) (Table S2). We consider it likely that these widespread sets of co-occurring proteins are foundations upon which lineage-specific metabolisms are based.

We conducted an unsupervised clustering of the genomes based on the presence/absence profiles of the 786 families considered to be widely distributed across the genomic dataset. As seen in Figure 2, the results clearly distinguish the CPR bacteria from other bacteria and archaea (Figure 3A). The interesting exception is the Dependentiae phylum (TM6), which is nested in the CPR group between Microgenomates and Parcubacteria (Figure 3B) although phylogenetic trees based on core genes clearly place Dependentiae outside of the CPR (Brown et al., 2015).

**Figure 3.**
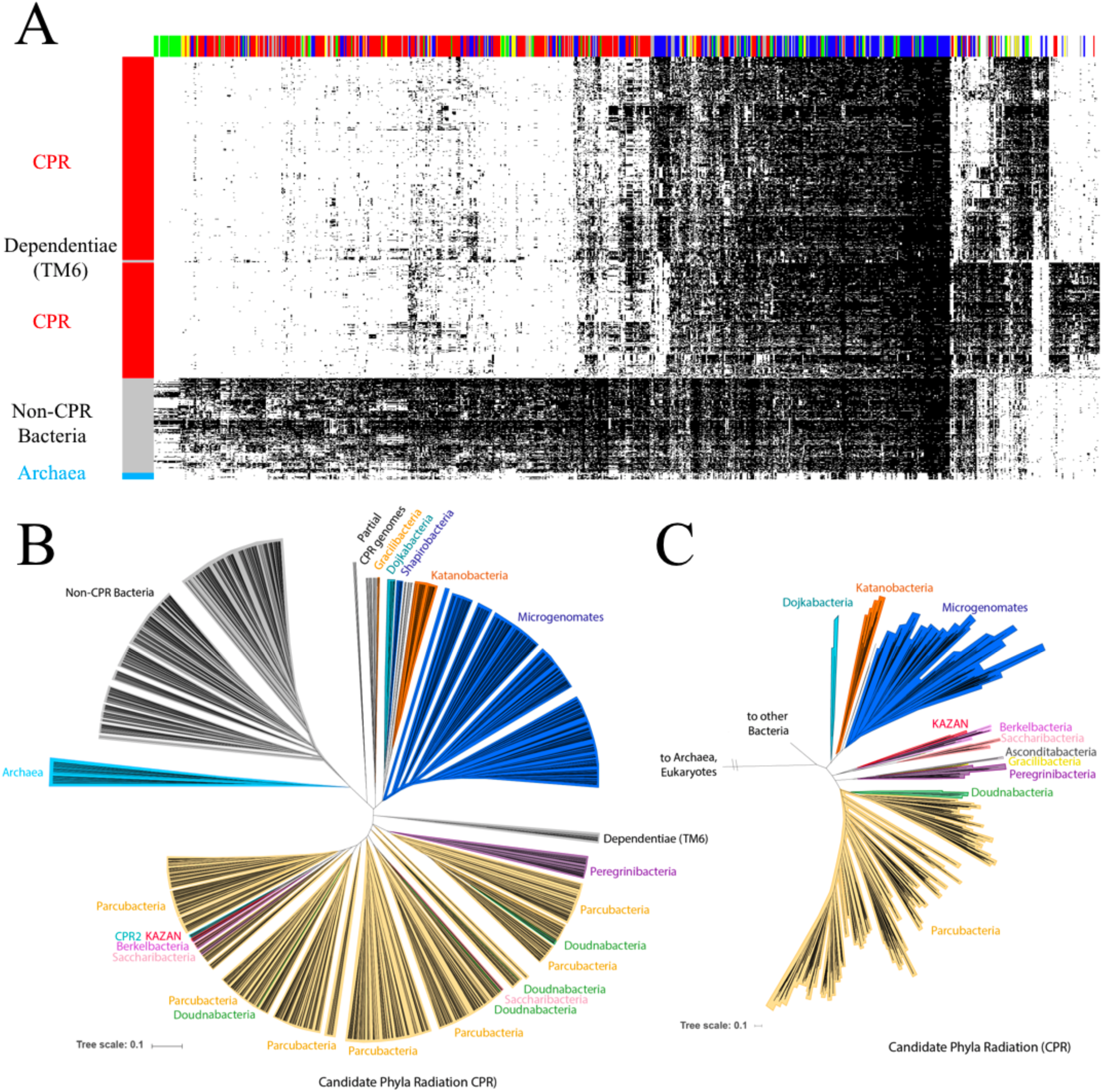
(A) The distribution of 786 widely distributed protein families (columns) in 2890 genomes (rows) from CPR bacteria (red), non-CPR bacteria (gray), and a few archaea (light blue) in a reference set with extensive sampling of genomes from metagenomes (thus includes sequences from many candidate phyla). Data are clustered based on the presence (black) / absence (white) profiles (Jaccard distance, complete linkage). Only near-complete and non-redundant genomes were used. The colored top bar corresponds to the functional category of families (Metabolism: red, Genetic Information Processing: blue, Cellular Processes: green, Environmental Information Processing: yellow, Organismal systems: orange, Unclassified: grey, Unknown: white). (B) Tree resulting from the hierarchical clustering of the genomes based on the distributions of proteins families in the panel A. (C) A phylogenetic tree of the CPR. Maximum-likelihood tree was calculated based on the concatenation of 14 ribosomal proteins (L2, L3, L4, L5, L6, L14, L15, L18, L22, L24, S3, S8, S17, and S19) using the PROTCATLG model.

Remarkably, within the CPR, genomes are clustered together based on the protein family distribution in patterns that are generally consistent with their phylogeny (Figures 3B and 3C). The Microgenomates and Parcubacteria superphyla form two distinct groups (Figure 3B), with Dojkabacteria (WS6) and Katanobacteria (WWE3) as sibling groups to Microgenomates and Doudnabacteria, Saccharibacteria, Berkelbacteria, Kazan and the Peregrinibacteria as sibling to, or nested in, Parcubacteria. When the hierarchical clustering pattern from the y-axis of Figure 3A is rendered in a radial tree format (Figure 3B) the correspondence between clusters based on the distribution of core protein families and phylogeny (Figure 3C) is particularly apparent.

The analysis present in Figures 2 and 3 used a genomic dataset that was notably enriched in CPR bacteria. To test whether the clear separation of CPR and non-CPR bacteria is an artifact of the choice of genomes, we created a second dataset of 2,729 of publicly available NCBI genomes sampled approximately at the level of one per genus (see Materials and Methods). The 786 protein families were identified in this dataset and arrayed using the same approach as in Figure 3A (Figure 4A). The diagram clearly separates CPR from non-CPR bacteria and from archaea. Thus, we conclude that the major subdivision within the first dataset was not due to our choice of genomes or the environments they came from. Importantly, this NCBI genome dataset includes many genomes from symbionts with reduced genomes (McCutcheon and Moran, 2012). In no case did these genomes place within the CPR.

**Figure 4.**
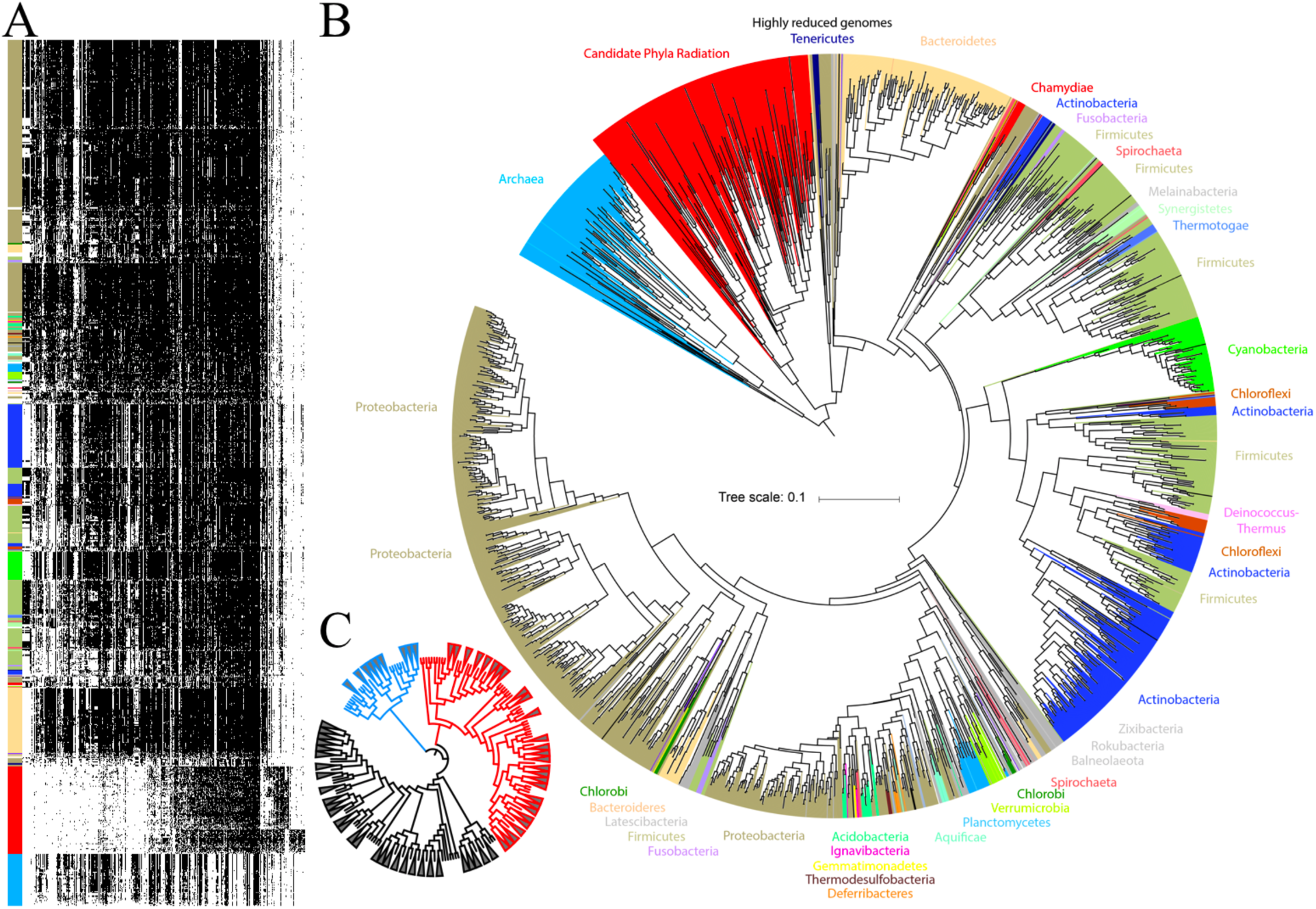
(A) The distribution of 786 widely distributed protein families (columns) in 2,616 near-complete and non-redundant genomes (rows) from a reference set with extensive sampling of genomes from non-CPR bacteria. Genomes are clustered based on the presence (black) / absence (white) profiles (Jaccard distance, complete linkage). The order of the families is the same as in Figure 3.A. (B) Tree resulting from the hierarchical clustering of the genomes based on the distributions of proteins families in the panel A. (C) The same tree with a collapsing of all branches that represented less than 25% of the maximum branch length (CPR are in red, Archaea in blue and non-CPR bacteria in grey).

From the hierarchical clustering of the genomes in Figure 4 we generated a tree representation analogous to that in Figure 3B (Figure 4B). Again, the correspondence between genome clusters based on protein family distribution and phylogeny is striking. For example, the 85 cyanobacterial genomes (bright green in Figure 4A) have a highly consistent pattern of presence/absence of core protein families and this is reflected in the comparatively short branch lengths in Figure 4B. In contrast, the branch lengths associated with the CPR bacteria are very long.

To evaluate branch length patterns through Figure 4B, we collapsed all branches that represented less than 0.25 of the maximum branch length (Figure 4C). In this rendering, the 99% of the Cyanobacteria collapse into a single wedge (84 out of 85). The CPR comprise 98 wedges, non-CPR bacteria 97 wedges and Archaea, 35 wedges. Notably, DPANN archaea cluster separately from other archaea, consistent with their phylogenetic separation in some analyses (Williams et al., 2017). The high representation of wedges of CPR relative to non-CPR bacteria is striking, given that CPR genomes represent only 11% of all genomes used in this analysis. We attribute this to high diversity in the subsets of core protein families present in genomes of organisms from across the CPR.

To test whether the protein clustering cutoffs strongly affected our results we performed another protein clustering without using cut-offs set during the HMM-HMM comparison (see Materials and Methods). This very inclusive procedure led to the definition of 3,555 clans (clusters of protein families that were identified in at least 5 distinct genomes). Despite merging of non-homologous proteins, we retrieved 537 clans that correspond to the 786 widespread families. Using this inclusive clustering, the CPR still separate from non-CPR bacteria and archaea in analyses that used both genome datasets (Figures S3 and S4). Thus, we conclude that, our results are robust regarding both genome selection (as tested using the NCBI genome dataset) and the protein clustering parameters.

### Biological capacities explain the singularity of CPR

To explore the reasons for the genetic distinction of CPR from non-CPR bacteria and archaea we divided the 786 protein families into three sets based on their abundances in CPR and in non-CPR Bacteria (Figure 5A). A set of 246 families are equally distributed across the Bacteria and 540 families are either depleted or enriched in CPR bacteria. The similarly distributed set contains families mostly involved in informational processes, primarily in translation (Table S2). The set of 453 families that are sparsely distributed in CPR yet very common in other Bacteria is enriched in metabolic functions (Fig. 3A and 5A). The remaining 87 protein families enriched in CPR and rare in non-CPR bacteria are discussed in detail below. Importantly, when these 87 protein families are removed from the set of widespread families and the analysis re-performed, the CPR still separate from all other bacteria both genome datasets (Figures S5 and S6).

**Figure 5.**
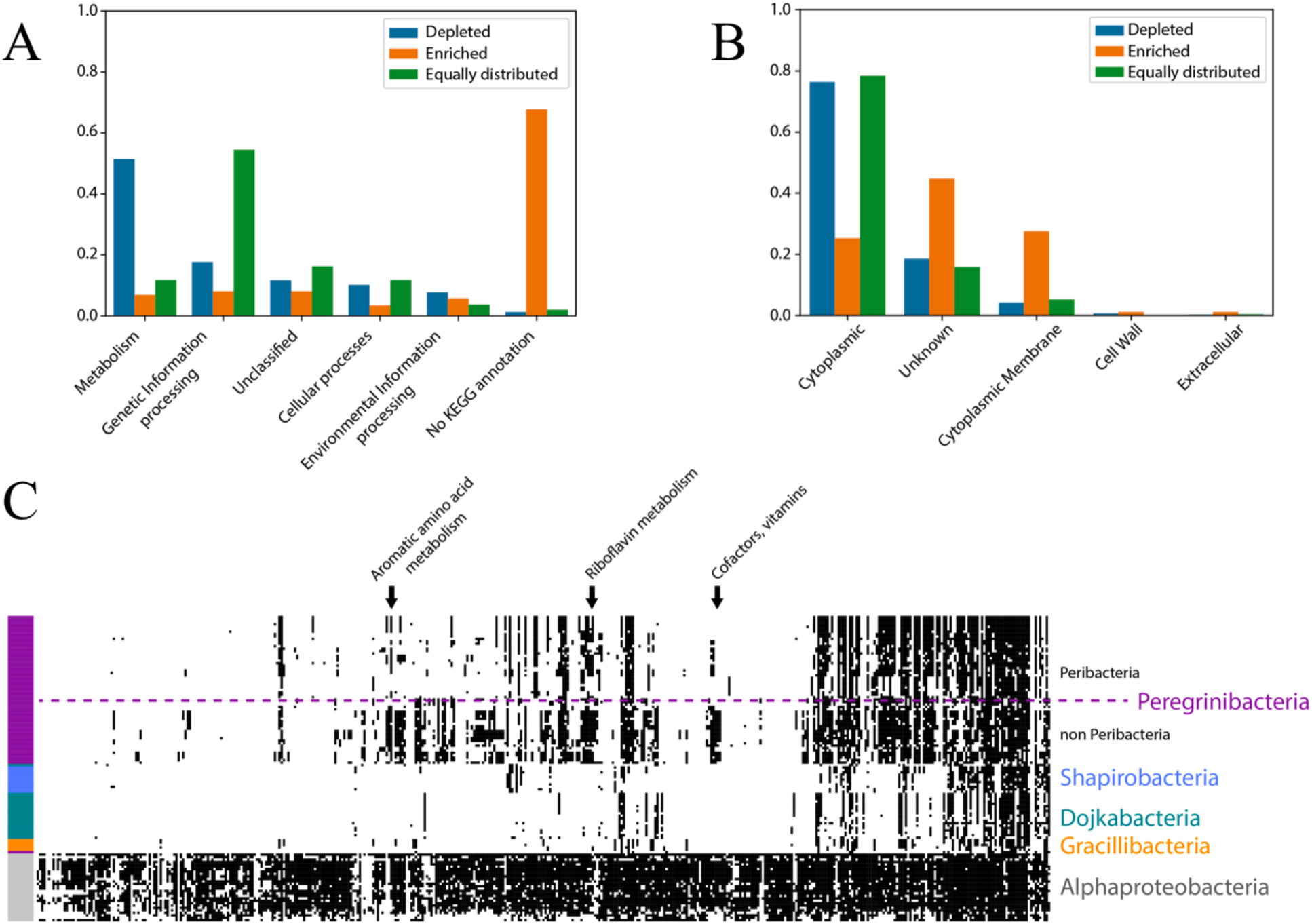
(A) Barplots of the distributions of the functional categories of the 786 protein families. (B) Barplots of the distributions of the cellular localizations of the 786 protein families. (C) Distribution of the 453 families that are depleted in CPR across 126 genomes from Peregrinibacteria (62), Shapirobacteria (11), Dojkabacteria (20), Gracilibacteria (5) and Alphaproteobacteria (28). The dashed line separates classes within the Peregrinibacteria. The order of the families and the genomes is the same as in Figure 3A.

Although the CPR are distinct from non-CPR bacteria due to their sparse metabolism and the presence of CPR-specific genes, they are not monolithic in terms of their metabolism (Figure 5C). For example, the genomes of the Peregrinibacteria encode far more metabolic families than Shapirobacteria, Dojkabacteria and Gracilibacteria. Despite the comparatively high metabolic gene inventory of the Peregrinibacteria, they have far fewer capacities than, for example, Alphaproteobacteria (Figure 5C).

### Many novel protein families enriched in CPR are linked to pili and cell-cell interactions

As noted above, 87 protein families are enriched in the CPR relative to non-CPR bacteria (Figure 6 and Table S2). Notably, Dependentiae genomes encode few or none of the 87 CPR-enriched families, consistent with their phylogenetic placement outside of the CPR (Figure 6). The majority of the 87 families has poor functional annotations (Figure 5A and Table S2). However, 51 families are comprised of proteins with at least one predicted transmembrane helix (Table S2), and many are predicted to have membrane or extracellular localizations (Figure 5B and Table S2). Fifteen have more than four transmembrane helices, and may be involved in transport (Table S2).

**Figure 6.**
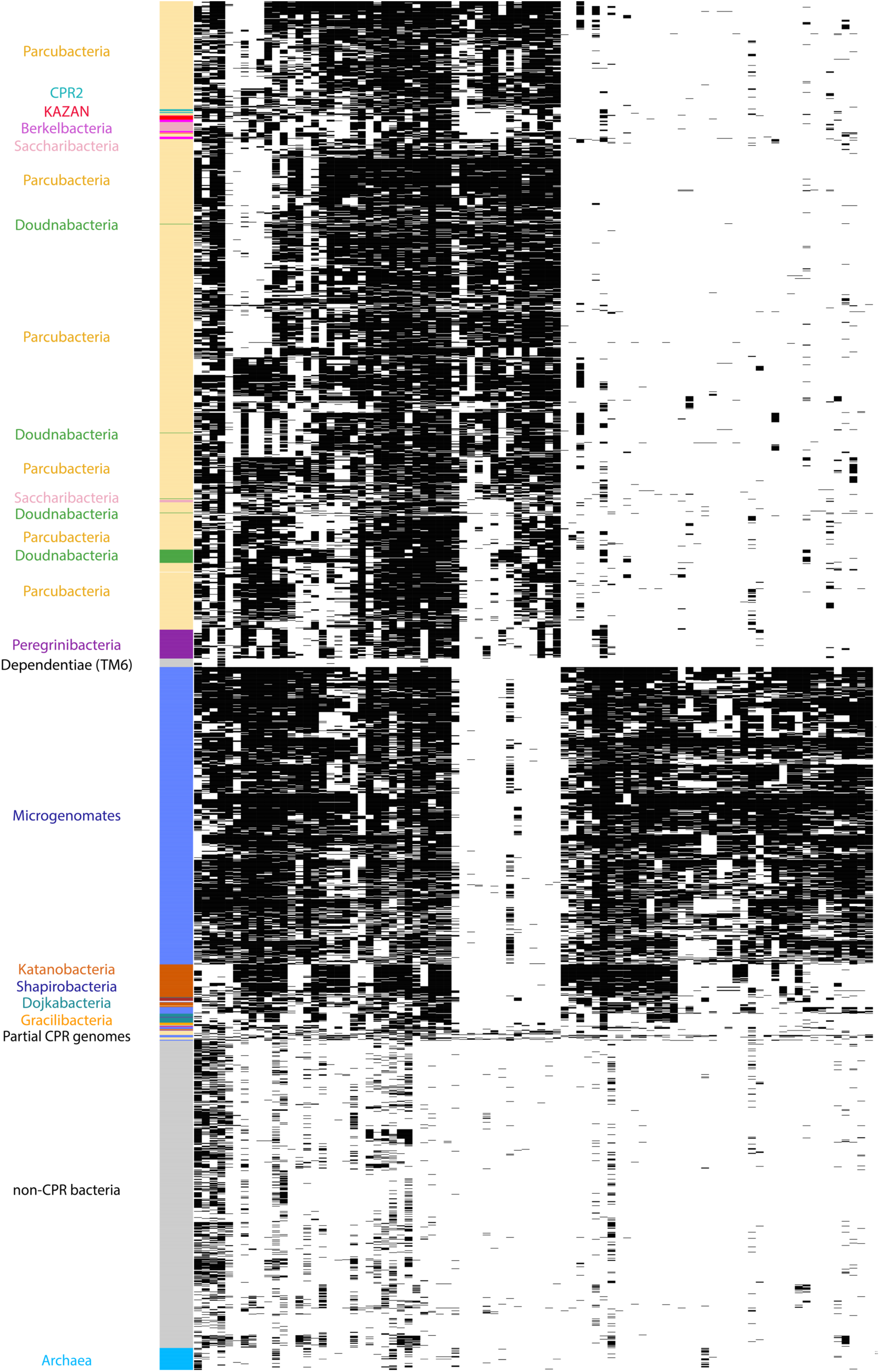
Distribution of the 87 families that are enriched in CPR relative to non-CPR Bacteria. The order of the families and the genomes is the same as in Figure 3A.

Interestingly, 28 of the 87 protein families are widespread in all CPR bacteria, the others are enriched in either Microgenomates (center, right side of Figure 6) or in Parcubacteria (center top in Figure 6). Those strongly associated with Microgenomates or Parcubacteria are primarily hypothetical proteins. However, 36 of 41 protein families enriched in Microgenomates are predicted to be non-cytoplasmic whereas 10 of 18 protein families enriched in Parcubacteria are predicted to be non-cytoplasmic (Table S2).

Given that most CPR are predicted to depend on externally-derived nucleic acids, it is anticipated that their cells are competent (Wrighton et al., 2016). This capacity is not widespread in non-CPR bacteria (Chen and Dubnau, 2004). We identified two families (3.6k.fam00166 and 3.6k.fam01878) annotated as ComEC (components of the DNA uptake machinery) among the 28 widespread CPR-enriched families (Table S2). Two other components of DNA uptake machinery, ComFC/comFA (3.6k.fam03513) and DprA (3.6k.fam01708), are also widespread in CPR but they are also widespread in non-CPR Bacteria. The ComEA component involved in DNA binding is only present in around one third of CPR genomes, suggesting that some CPR may possess an alternative mechanism for DNA binding.

In competent bacteria, a correlation has been shown between the ability to take up exogenous DNA and the presence of pili on the cell surface (Stone and Kwaik, 1999). We found that three enriched and widespread families in CPR are divergent pilin proteins, the subunits of pili. These typically have a single transmembrane domain in their first 50 amino-acids (families 3.6k.fam00087, 3.6k.fam00099, 3.6k.fam00143) (Giltner et al., 2012). These pilin proteins are part a type IV pili (T4P) system that includes other components that are encoded in the genomes of CPR bacteria but also present in non-CPR Bacteria (Melville and Craig, 2013). These components comprise the ATPase assembly PilB (3.6k.fam04160), the ATPase twitching motility PilT (3.6k.fam00968), the membrane platform PilC (3.6k.fam00031), the prepilin peptidase PilD (3.6k.fam02501) and finally the Gspl domain PilM (3.6k.fam00032). All of these components co-localize in several CPR genomes. Importantly we did not find the PilQ component which is required to extrude the pilus filament across the outer membrane of gram negative Bacteria (Chen and Dubnau, 2004), consistent with the microscopy observations that suggest CPR do not have a gram negative cell envelope (Luef et al., 2015).

Full-length type IV pilin precursors are secreted by the Sec pathway in unfolded states in gram positive bacteria (Giltner et al., 2012). A thiol-disulfide oxidoreductase (3.6k.fam00998) is one of the protein families enriched in CPR bacteria and may be involved in ensuring correct folding of the pilins. These proteins show similarity to membrane-bound oxidoreductase MdbA, which is found in the gram positive *Actinomyces oris* (Reardon-Robinson et al., 2015a) and *Corynebacterium diphtheriae* (Reardon-Robinson et al., 2015b). In these organisms, MdbA catalyzes disulfide bond formation in secreted proteins, a reaction that is important for protein stability and function (Reardon-Robinson and Ton-That, 2016). In *Actinomyces oris*, one of these secreted proteins is the FimA pilin (Reardon-Robinson et al., 2015a). Similarly to MdbA, 59% of the proteins from the family 3.6k.fam00998 are predicted to be anchored in the cell wall and the catalytic CxxC motif required for disulfide bond formation is conserved in 3223 of 3496 (92%) CPR proteins. Interestingly, the family of proteins most frequently found in genomes adjacent to proteins of 3.6k.fam00998 in CPR bacteria is the Vitamin K epoxide reductase (VKOR, adjacent in 76 CPR genomes). VKOR re-oxidizes MdbA in *Actinomyces oris* (Luong et al., 2017).

Another interesting protein family with one or more transmembrane segments (3.6k.fam08817) that is widespread and almost unique to CPR organisms possesses a CAP domain (Cysteine-rich secretory proteins, Antigen 5, and Pathogenesis-related 1 proteins). The CAP domains are found in diverse extracellular proteins of bacteria and eukaryotes and have a wide range of physiological activities including fungal virulence, cellular defense and immune evasion (Gibbs et al., 2008). Some members of this family are endopeptidases, some are transglycosylases or have non-enzymatic roles. Many other CPR-enriched families were identified but lack confident functional predictions (Table S2).

## Discussion

Genome-resolved metagenomics studies have greatly expanded our understanding of microbial life, particularly through discovery of new bacterial lineages (candidate phyla). Lacking have been studies that investigate these genomes from the perspective of the diversity and distribution patterns of homologous proteins. To begin comparing protein sequence inventories, we clustered the amino acid sequences into families that approximate homologous groups. These families serve as a common language that enables comparison of gene inventories within and among lineages. Strikingly, the combinations of protein families associated with widespread biological capacities separate the CPR from all other bacteria. In other words, the pattern of presence/absence of relatively widely distributed protein families highlights a major dichotomy within Domain Bacteria that corresponds almost exactly to the subdivide inferred based on phylogenetic analyses (both rRNA and concatenations of ribosomal proteins) (Hug et al., 2016).

One potential explanation for the phylogenetic separation of CPR bacteria from other bacteria is that the radiation is comprised of fast-evolving symbionts. If this was the case, then we might predict that other fast evolving bacterial symbionts (e.g., those associated with insects, Chlamydia, Mollicutes and other bacteria with highly reduced genomes) would cluster phylogenetically with the CPR. This is not the case. The separation based on presence/absence patterns persists even when the analysis is redone without inclusion of the CPR-enriched families. Perhaps even more importantly, CPR bacteria separate from other bacteria, including other bacterial symbionts, based on essentially CPR-specific genes. The near ubiquity of 87 protein families across the CPR radiation is most readily explained by early acquisition at the time of the origin of CPR, with persistence via vertical inheritance.

Based on the functional predictions, the protein families that are highly enriched in CPR and rare or absent in other bacteria may be important for interaction between CPR and their hosts. Among them, the type IV pili may be central to CPR associations with other organisms. These molecular machines confer a broad range of functions from locomotion, adherence to host cells, DNA uptake, protein secretion and environmental sensing (Maier and Wong, 2015). Prominent among this set are three novel pilin protein families that may have evolved in CPR bacteria. Notably, several other groups of CPR-enriched genes are also predicted to function in DNA uptake and maintenance of pilin structure. Given the overall small genome size, these findings reinforce the conclusion that genes for organism-organism interaction are central to the lifestyles of CPR bacteria.

Similar to CPR bacteria, phylum-level groups of non-CPR bacteria also have unique sets of core genes. For example, Cyanobacteria group together tightly despite the fact that genes for the physiological trait that unites them, oxygenic photosynthesis, were not included in the protein family analysis. This may be a reflection of the requirement for a specific combination of core protein families that comprise a platform that is consistent with photosynthetic lifestyles. Similar explanations are less easily identified for the general correspondence of phylogeny and conserved protein family sets in other major groups. However, a general explanation may be that once an innovation that gave rise to a lineage occurred, strong selection maintained the core protein family platform set that supports it.

Within the CPR we identified many clusters of bacteria that share similar core metabolic platforms. Some CPR bacterial phyla have extensive biosynthetic capacities whereas others have minimal sets of core protein families. This may indicate extensive gene loss in some groups. Given the overall phylum-level consistency of the protein family sets, we suspect that major genome reduction events were ancient.

Looking across the entire analysis, the broad consistency in combinations of core protein families within lineages strongly suggests that the distribution of these families is primarily the result of vertical inheritance. Specifically, the patterns of protein family distribution reproduce the subdivision of Bacteria from Archaea and essentially recapitulate many phylum and sub-phylum groupings. Collapsing the branches in the cladogram formed from the hierarchical clustering of protein families revealed enormous branch length in the CPR. We interpret this to indicate huge variation in the sets of core protein families across the CPR (Figure 4C). This diversity in the core protein family platform may have arisen because distinct types of symbiotic associations select for larger or lesser requirement for core biosynthetic capacities in the symbiont. In this case, major divergences within the CPR, essentially the rise of new CPR phyla, may have been stimulated by evolutionary innovations that generated new lineages of potential bacterial hosts.

The relative magnitude of diversity of distinct core gene sets in the CPR compared to non-CPR bacteria is consistent in scale with the relative magnitude of phylogenetic diversity of these groups, as rendered in ribosomal RNA and protein trees (Hug et al., 2016). This suggests the enormous biological importance of the CPR, regardless of the extent to which their phylogenetic and core metabolic diversity is a reflection of gene loss, rapid evolution or ancient origin within the bacterial domain.

## Materials and Methods

### Datasets construction

The initial dataset contains 3,598 prokaryotic genomes (5,061,957 proteins) that were retrieved from 4 published datasets (Anantharaman et al., 2016; Brown et al., 2015; Castelle et al., 2015; Probst et al., 2017). The dataset encompasses 2,321 CPR (1,953,651 proteins); 1,198 non-CPR-Bacteria (3,018,597 proteins) and 79 Archaea (89,709 proteins) (Table S3). The second “NCBI” dataset contains 2,729 genomes (8,425,478 proteins). Genome were chosen based on the taxonomy provided by the NCBI. Briefly, for each prokaryotic phylum, one genome per genus was randomly selected from the NCBI genome database (last access on December, 2017). Some genomes do not have genus assignment although they have a phylum assignment. In those cases, 5 genomes per phylum were randomly selected. Refseq were preferred to non-refseq genomes as these are generally better annotated. The NCBI dataset encompasses 282 CPR (217,728 proteins); 2,278 non-CPR-Bacteria (7,811,207 proteins) and 169 Archaea (396,543 proteins) (Table S3).

### Protein clustering

Proteins with ≥ 90% identity were clustered using CD-HIT (Fu et al., 2012) to remove nearly identical proteins and protein fragments, and representatives of each cluster where used in downstream protein clustering. All vs. all local searches were performed using usearch (Edgar, 2010) with -ublast and -evalue 0.0001 parameters, and the bit-score was used for MCL (Markov CLustering algorithm) (Enright et al., 2002), with 2.0 as the inflation parameter. Each of the resulting clusters that included at least 5 representative proteins was sub-clustered based on global percent identity. This was achieved by performing an all vs. all search within the members of the cluster with usearch64-search_global and performing MCL based on percent identity (with-I 2.0). The resulting sub-clusters were defined as sub-families. In order to test for highly similar sub-families, we grouped sub-families into protein families as follows. The proteins of each sub-family were aligned using MAFFT (Katoh and Standley, 2013), and from the alignments HHM models were built using the HHpred suite (Soding, 2005). The sub-families were then compared to each other using hhblits (Remmert et al., 2011) from the HHpred suite (with parameters -v 0 -p 50 -z 4 -Z 32000 -B 0 -b 0). For sub-families with probability scores of ≥ 99% and coverage ≥ 0.75, a similarity score (probability ⨉ coverage) was used in the final MCL (-I 2.0). These clusters were defined as the protein families, after adding to each representative highly similar proteins that were removed in the first CD-HIT step. The clans were defined by clustering all sub-families using MCL based on the probability ⨉ coverage score (without using the cutoffs of probability and coverage used for the protein family clustering).

### Selection of widespread families

Examining the distribution of the protein families across the genomes, a clear modular organization emerged (Figure 2. Panel A). We used the Louvain algorithm (Blondel et al., 2008) to detect modules of proteins that share similar patterns of presence/absence across the genomes. Briefly, The Louvain algorithm seeks a partition of a network that maximizes the modularity index Q. The algorithm was performed on a weighted network that was built by connecting family nodes sharing a Jaccard index greater than 0.5. For each pair of protein families, the Jaccard index was calculated based on their profiles of presence/absence across the genomes. The 0.5 threshold was empirically chosen because it defined 3 distinct modules for widespread proteins in Archaea, non-CPR-Bacteria and Bacteria (see Figure 2A) whereas lower thresholds merged families having distinct presence/absence patterns across the genomes. This procedure defined modules with more than 10 proteins.

For each module, the genomes with that module were identified and their phylum affiliations determined. Briefly, for each module, the median number of genomes per family (*m*) was calculated. The *m* genomes that contains the biggest number of proteins were retained; their phyla distribution defines the taxonomic assignment of the module.

### Genome completeness assessment and de-replication

Genome completeness and contamination were estimated based on the presence of single-copy genes (SCGs) as described in (Anantharaman et al., 2016). For CPR, 43 universal SCGs were used, following (Anantharaman et al., 2016). In non-CPR bacteria, genome completeness was estimated using 51 SCGs, following (Anantharaman et al., 2016). For archaea, 38 SCGs were used, following (Anantharaman et al., 2016). Genomes with completeness > 70% and contamination < 10% (based on duplicated copies of the SCGs) were considered as near-complete genomes. Genomes were de-replicated using dRep (Olm et al., 2017) (version v2.0.5 with ANI > 95%). The most complete genome per cluster was used in downstream analyses.

### Functional annotation

Protein sequences were functionally annotated based on the accession of their best Hmmsearch match (version 3.1) (E-value cut-off 0.001) (Eddy, 1998) against an HMM database constructed based on ortholog groups defined by the KEGG (Kanehisa et al., 2016) (downloaded on June 10, 2015). Domains were predicted using the same Hmmsearch procedure against the Pfam database (version 31.0) (Punta et al., 2012). The domain architecture of each protein sequence was predicted using the DAMA software (default parameters) (Bernardes et al., 2016). SIGNALP (version 4.1) (parameters: -f short -t gram+) (Petersen et al., 2011) and PSORT (version 3.0) (parameters: --long --positive) (Peabody et al., 2016) were used to predict the putative cellular localization of the proteins. Prediction of transmembrane helices in proteins was performed using TMHMM (version 2.0) (default parameters) (Krogh et al., 2001).

### Enrichment analysis

Enrichment/depletion of protein families was calculated based on the frequency of the computed protein families in UniProt’s Reference Proteome database (downloaded April 17, 2017). First, a database of all HMMs of the sub-families was used to identify members of each sub-family in the Reference Proteomes from CPR and non-CPR bacteria. Additionally, HMM representing 16 single copy ribosomal genes were used to identify those proteins. The enrichment of each family in CPR vs. non-CPR bacteria was then computed using a Fisher exact test, in which the expected values were the count of single copy ribosomal genes in CPR and non-CPR, and the observed values were the counts of members of each protein families in CPR and non-CPR bacteria. Families were considered enriched or depleted if their p-values, after correction for false detection rate, were significant (< 10-5) and if their odd ratio were > 2. The remaining families were assigned as equally distributed.

## Acknowledgments

Support was provided by grants from the Lawrence Berkeley National Laboratory’s Genomes-to-Watershed Scientific Focus Area. The U.S. Department of Energy (DOE), Office of Science, and Office of Biological and Environmental Research funded the work under contract DE-AC02-05CH11231 and the DOE carbon cycling program DOE-SC10010566, the Innovative Genomics Institute at Berkeley and the Chan Zuckerberg Biohub. D.B. was supported by a long-term EMBO fellowship

## Author Contributions

RM, DB, CC and JB designed the analysis. DB assembled the initial dataset and performed the protein clustering. RM detected the widespread families and created the binary matrix. RM performed the functional analysis. CC performed the phylogenetic tree. All authors contributed to the analysis of the data and the interpretation of the results. All authors wrote the manuscript. All authors read and approved the final manuscript.

## Declaration of Interests

The authors declare no competing interests.

## Deposited data

All genomes used in the analysis are publicly available (see Table S3). The fasta sequences of the 786 families and the binary matrix used to create the figures 2, 3 and 4 are available at https://doi.org/10.6084/m9.figshare.6296987.v1.

## Supplementary Materials

**Figure S1.**
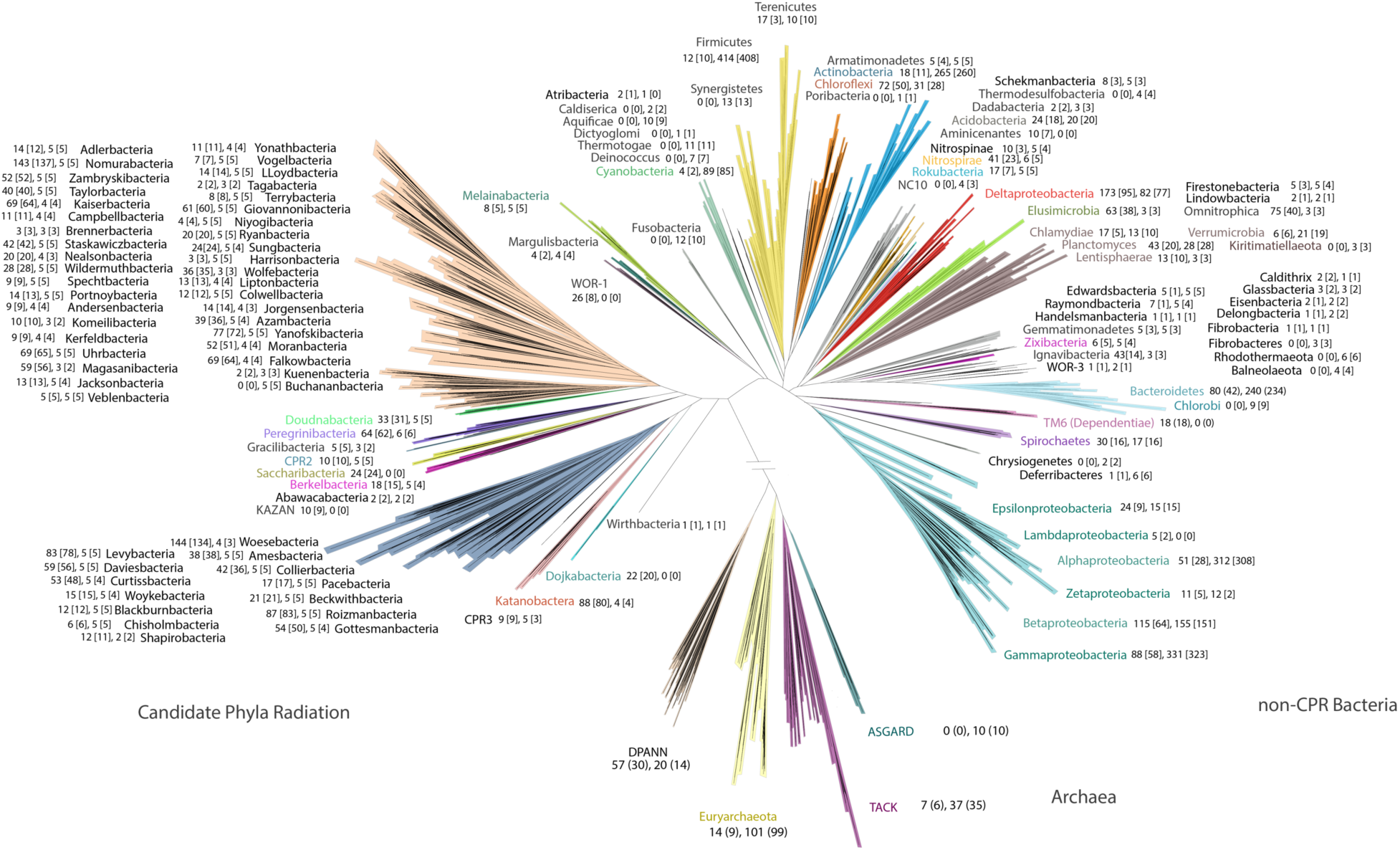
Tree illustrating phylogenetic sampling used in this study (the diagram is based on a tree published recently in (Castelle and Banfield, 2018)). For each phylum, the number of genomes and near-complete genomes (square brackets) is reported for the two datasets.

**Figure S2.**
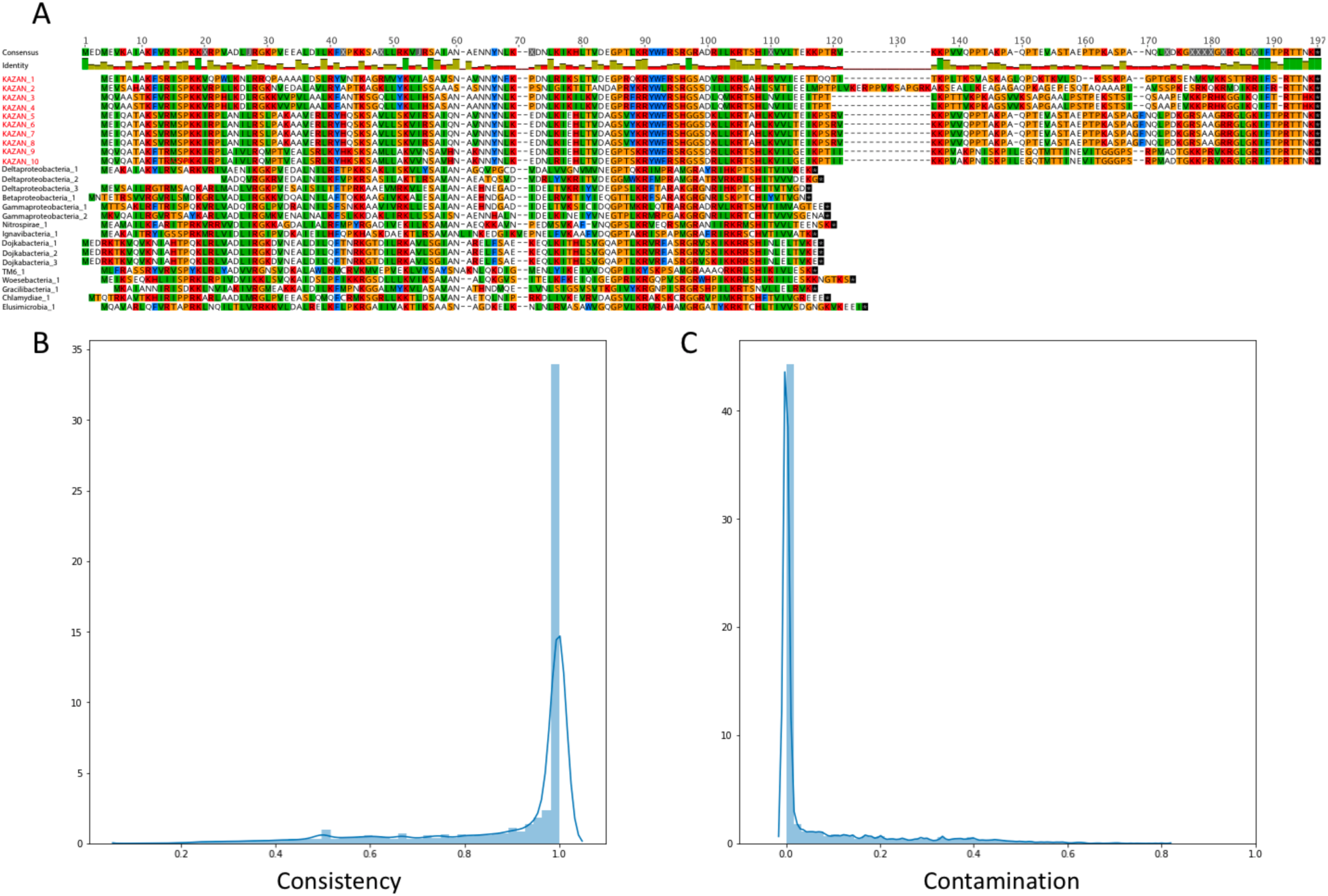
Quality assessment of the protein clustering. A. A multiple sequences alignment (MSA) of the 10 protein sequences from the family 3.6k.fam10722 (names in red) and 16 proteins sequences from the family 3.6k.fam00371 (names in black). The MSA highlights the extension of the C-terminal region of proteins from the family 3.6k.fam10722. B. Histogram of the ratios of the most abundant KEGG accessions that are present in the most abundant families. C. Histogram of the ratios of admixture for each family.

**Figure S3.**
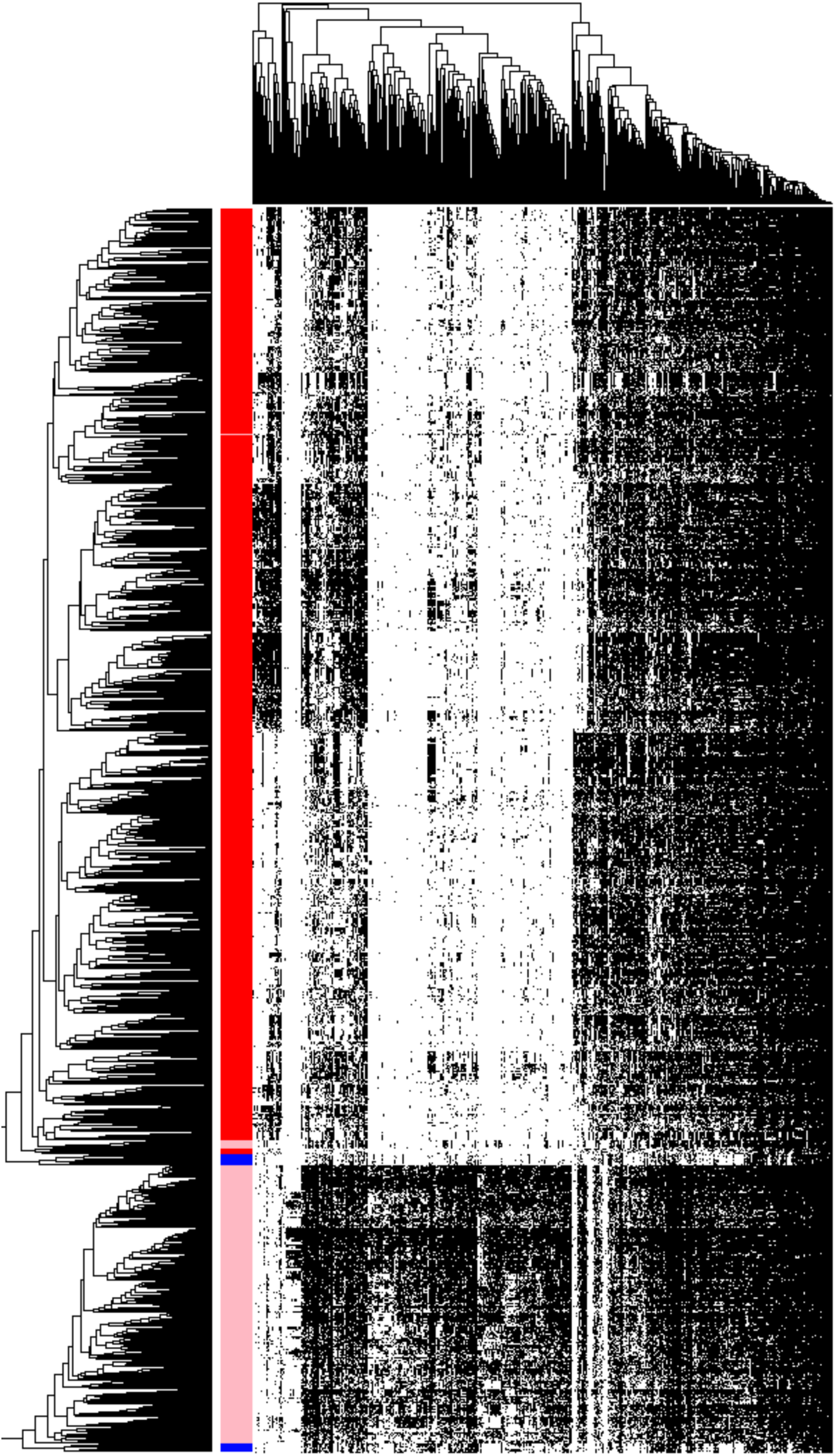
The distribution of widely distributed 537 protein clans (columns) in 2890 genomes (rows) from CPR bacteria (red), non-CPR bacteria (pink), and a few archaea (blue) in a reference set with extensive sampling of genomes from metagenomes (thus includes sequences from many candidate phyla). Data are clustered based on the presence (black) / absence (white) profiles (Jaccard distance, complete linkage). Only near-complete and non-redundant genomes were showed. The non-CPR bacteria genomes (in pink) that are nested in the CPR (in red) correspond to the dependentiae (TM6) genomes.

**Figure S4.**
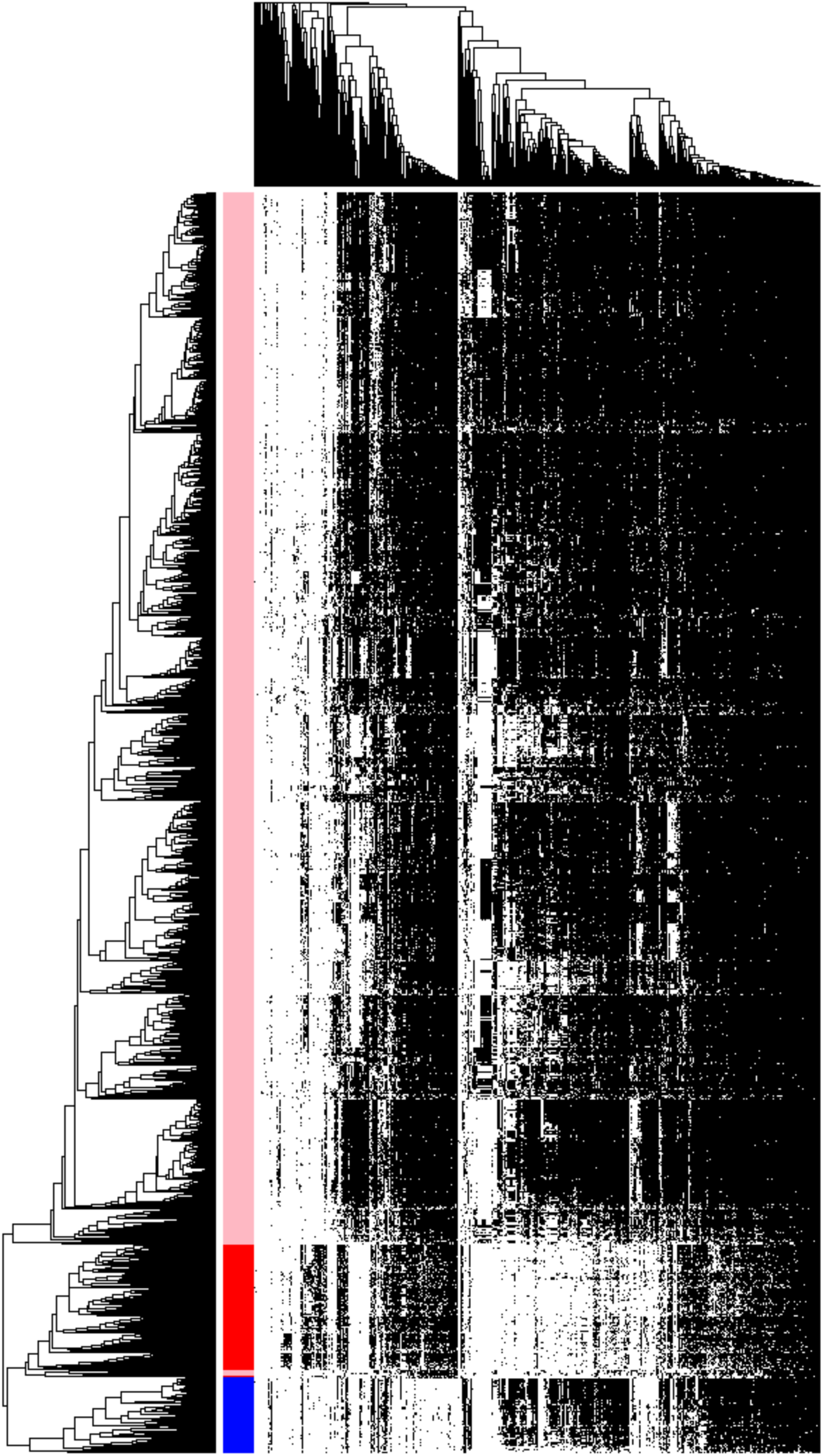
The distribution of 537 widely distributed protein clans (columns) in 2,616 near-complete and non-redundant genomes (rows) from a reference set with extensive sampling of genomes from non-CPR bacteria (pink). Genomes are clustered based on the presence (black) / absence (white) profiles (Jaccard distance, complete linkage). CPR bacteria are colored in red, the Archaea are colored in blue.

**Figure S5.**
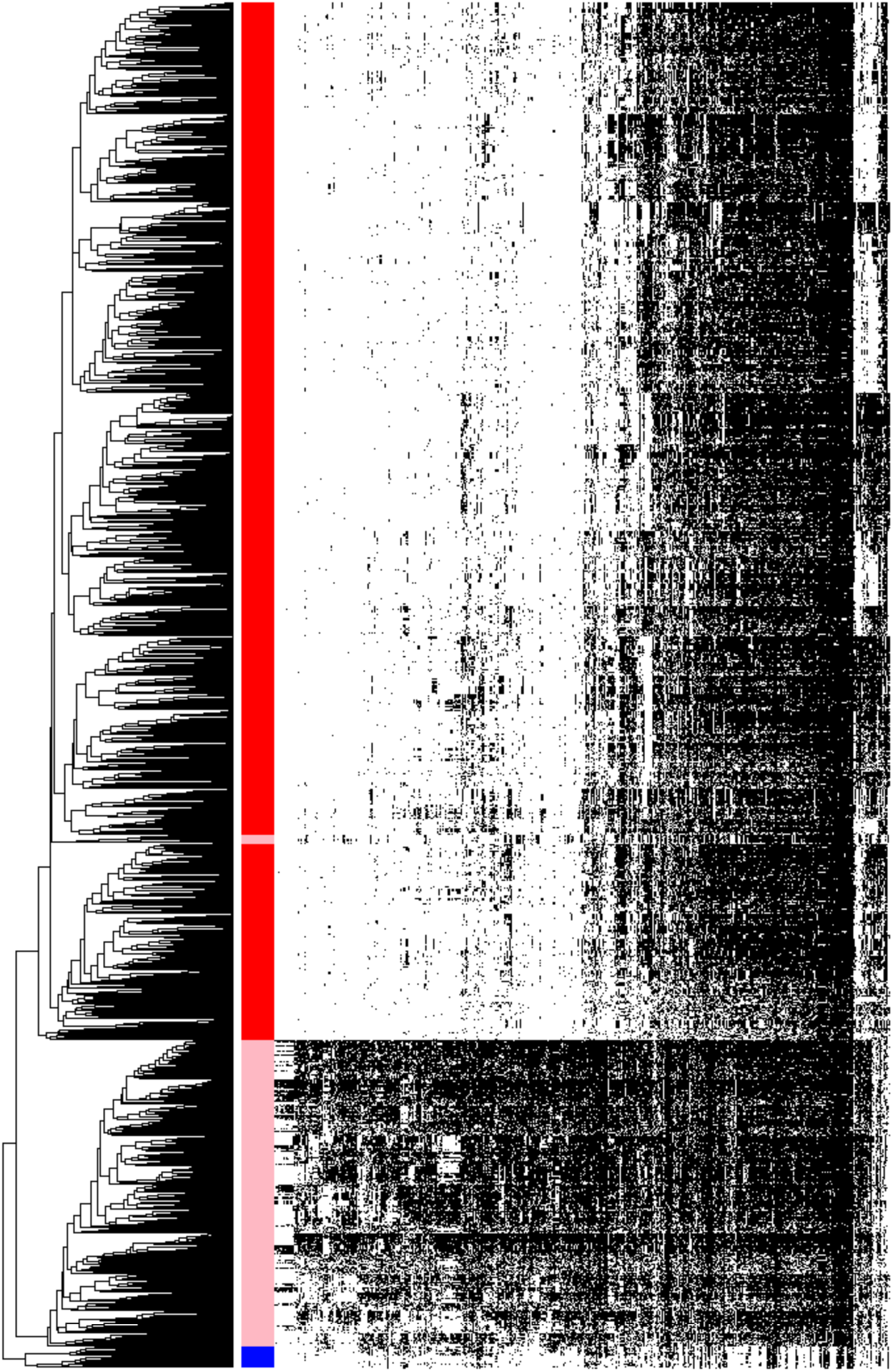
The distribution of 699 widely distributed protein families (columns) in 2890 genomes (rows) from CPR bacteria (red), non-CPR bacteria (pink), and a few archaea (blue) in a reference set with extensive sampling of genomes from metagenomes (thus includes sequences from many candidate phyla). The 87 families that are enriched in CPR relative to non-CPR bacteria were not considered in this analysis. Genomes are clustered based on the presence (black) / absence (white) profiles (Jaccard distance, complete linkage). Only near-complete and non-redundant genomes were showed. The order of the families is the same as in Figure 2A. The non-CPR bacteria genomes (jn pink) that are nested in the CPR (in red) correspond to the dependentiae (TM6) genomes.

**Figure S6.**
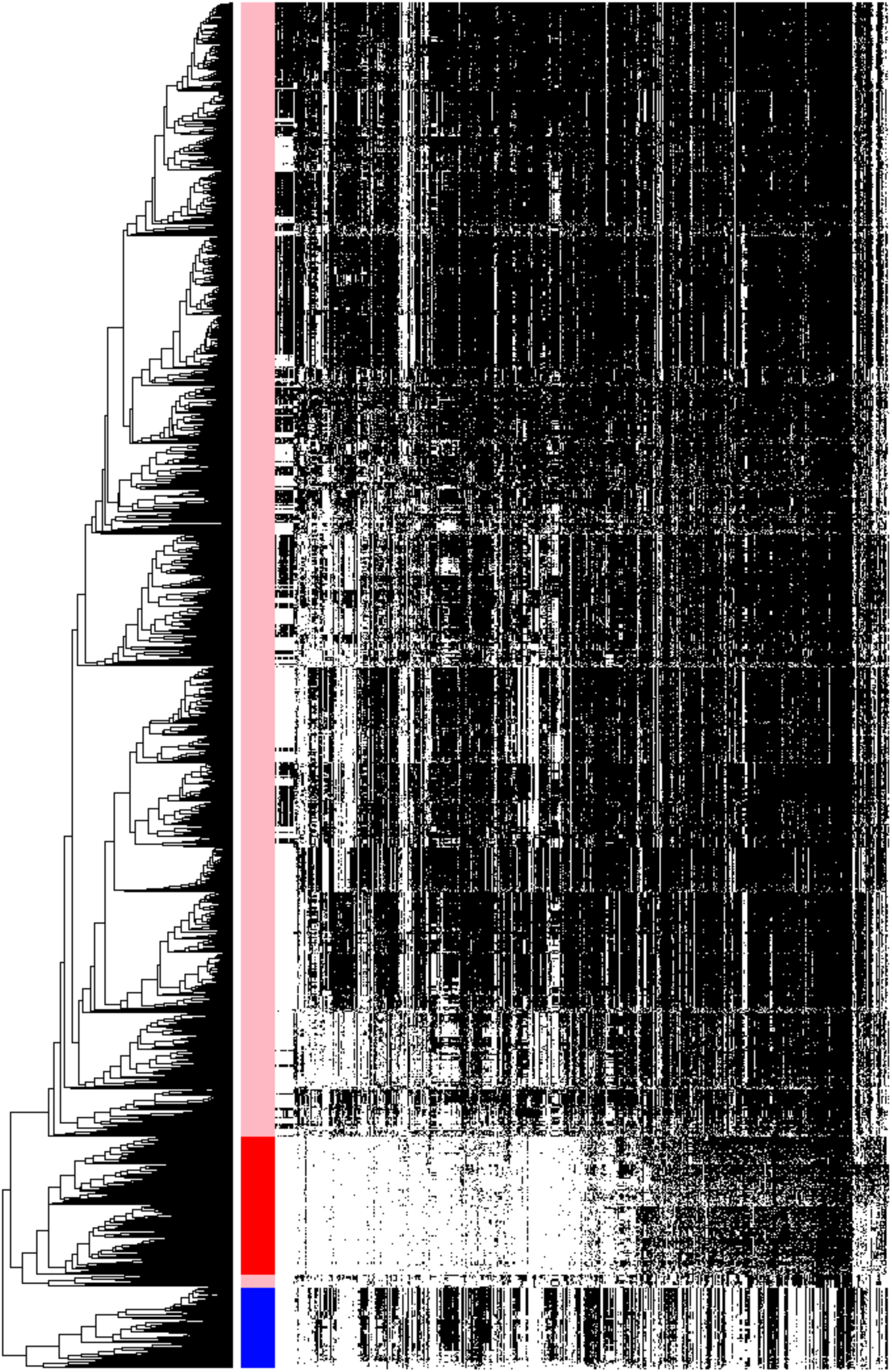
Tree resulting from the hierarchical clustering of the genomes based on the distributions of 699 widely distributed protein families in 2,616 near-complete and non-redundant genomes from a reference set with extensive sampling of genomes from non-CPR bacteria (pink). The 87 families that are enriched in CPR relative to non-CPR bacteria were not considered in this analysis. Genomes are clustered based on the presence (black) / absence (white) profiles (Jaccard distance, complete linkage). The order of the families is the same as in Figure 2A.

**Table S1.** Quality control of the protein clustering based on the 16 ribosomal proteins (RP). The rational is because those proteins are highly conserved, we expect to have each of them into a single cluster (i.e. one family per type of RP). This table summarized the results, each line corresponds to one RP protein. The proteins annotated as one of the 16 RP were retrieved using the corresponding Pfam accessions (PFAM accessions column). For 10 of them, all proteins cluster into one single family (# families column). The 6 remaining RP clusters in several families however there is always one family that contains the majority of the proteins (# proteins column).

| RP | Pfam accessions | # families | # proteins | Family Accession | # unclustered proteins |
| --- | --- | --- | --- | --- | --- |
| RPL2 | PF00181+PF03947 | 2 | 3497<br>54 | 3.6k.fam07672<br>3.6k.fam04518 | 3 |
| RPL3 | PF00297 | 1 | 3434 | 3.6k.fam03114 | 5 |
| RPL4 | PF00573 | 1 | 3455 | 3.6k.fam01116 | 30 |
| RPL5 | PF00281+PF00673 | 2 | 3462<br>70 | 3.6k.fam16812<br>3.6k.fam03533 | 17 |
| RPL6 | PF00347+PF00347 | 1 | 3509 | 3.6k.fam02196 | 5 |
| RPL14 | PF00238 | 1 | 3529 | 3.6k.fam21706 | 6 |
| RPL15 | PF00828 | 2 | 71<br>3362 | 3.6k.fam03860<br>3.6k.fam04045 | 2 |
| RPL16 | PF00252 | 2 | 53<br>3461 | 3.6k.fam06774<br>3.6k.fam08297 | 13 |
| RPL18 | PF00861 | 1 | 3393 | 3.6k.fam20082 | 2 |
| RPL22 | PF00237 | 6 | 3300<br>23<br>8<br>32<br>10<br>16 | 3.6k.fam00371<br>3.6k.fam10720<br>3.6k.fam13371<br>3.6k.fam02454<br>3.6k.fam10722<br>3.6k.fam10726 | 87 |
| RPL24 | PF17136 | 1 | 3408 | 3.6k.fam11163 | 11 |
| RPS3 | PF07650+PF00189 | 2 | 6<br>3535 | 3.6k.fam21121<br>3.6k.fam04284 | 27 |
| RPS8 | PF00410 | 1 | 3415 | 3.6k.fam02173 | 71 |
| RPS10 | PF00338 | 1 | 3268 | 3.6k.fam04313 | 5 |
| RPS17 | PF00366 | 1 | 3498 | 3.6k.fam02721 | 24 |
| RPS19 | PF00203 | 1 | 3485 | 3.6k.fam13933 | 0 |

Table S2. Annotation of the 786 widespread families. Column A: family accession. Column B: number of proteins in the family. Column C: median length of the proteins. Column D: ratio of proteins predicted to contain a signal peptide. Column E: median number of predicted transmembrane helix per protein. Column F: predicted cellular localization according to Psort. Column G: ggkbase annotation. Column H: domain architecture reported by Pfam. Columns I, J, K, L, M: KEGG annotations. Column N: number of CPR genomes that carry the family. Column O: number of non-CPR bacterial genomes that carry the family. Column P: number of DPANN genomes that carry the family. Column Q: number of non-DPANN archaeal genomes that carry the family. Column R: Abundancy category of the family in the CPR relative to non-CPR bacteria (depleted, enriched or equally distributed).

Table S3. List of the prokaryotic genomes we used in the comparative analysis. For each genome, the taxonomy, accession and the levels of completeness and contamination are provided.

